# Lipid and supramolecular structural protein signatures distinguish structurally colored from pigmented wing regions in the butterfly *Parides eurimedes mylotes*

**DOI:** 10.64898/2026.11.26.769165

**Authors:** Praveen Kumar Jaya Balaji, Colin Volet, Pierre-Marie Allard, Michael Stumpe, Maria Saba, Jorge Larios, Ullrich Steiner, Viola Vogler-Neuling

**Affiliations:** Adolphe Merkle Institute, University of Fribourg, Chemin des Verdiers 4, 1700 Fribourg, Switzerland; Department of Biology, University of Fribourg, Chemin du Musée 10, 1700 Fribourg, Switzerland; Metabolomics and proteomics platform, University of Fribourg, Chemin du Musée 10, 1700 Fribourg, Switzerland; National Center of Competence in Research Bio-inspired Materials, Chemin des Verdiers 4, 1700 Fribourg, Switzerland

## Abstract

Butterfly colouration arises from pigments and hierarchical nano- and microstructures, yet its molecular building blocks remain poorly resolved. Here, we developed an integrated region-specific lipidomics and proteomics workflow to identify the molecular determinants across wing regions of *Parides eurimedes mylotes*. Lipidomics revealed pronounced differences between photonic and pigmented regions, with the photonic region (Bragg stack) showing distinct diacylglycerol and triacylglycerol signatures. Canthaxanthin and lutein were identified as the yellow and red pigments. Proteomics revealed cytoskeletal, actin, and myosin proteins in both photonic and pigmented regions. The green photonic patch was additionally enriched in supramolecular polymers, fibres and complexes, contractile fibres, microtubules, sarcomeres and myofibrils, as well as proteins involved in lipid metabolism and vesicle-mediated transport. This data reveals coordinated spatial differences in lipid composition and protein profiles between pigmented and photonic regions, providing a biochemical dimension to photonic nanostructure formation and their biomimetic recreation.

---

Insects have evolved an extraordinary diversity of functional nanostructures that mediate adhesion, loco-motion, hydrophobicity, sensing and actuation [1]. These nanostructures offer considerable technological potential. Whereas biological systems assemble biodegradable materials with nanoscale precision under ambient conditions, comparable synthetic fabrication often relies on costly, energy-intensive, and defect-prone processes that remain difficult to scale to complex three-dimensional nanostructures. Rather than merely mimicking biological designs, identifying their molecular building blocks and elucidating the mechanisms by which they assemble could enable cell-free nanobiotechnological platforms for manufacturing advanced functional and hierarchical nanomaterials.

One prominent example is structural colour, which produces some of the most brilliant colours found in nature. These colours arise from the constructive interference of light within nanostructures exhibiting periodicities of several hundred nanometres [2]. Structural colouration frequently acts in combination with pigments, which can enhance or modulate the photonic response. Among the most remarkable three-dimensional photonic architectures is the gyroid geometry found in butterfly wing scales [3]. These nanostructures develop within the lumen of scales derived from epithelial cells [4]. Developmental studies of gyroid structures in *Papilionidae* and *Lycaenidae* have proposed that these nanostructures arise from membrane-defined templates within developing scale cells. In this model, a membrane-rich double gyroid is transformed into a single gyroid through the selective deposition of chitin [5]. More recently, the gyroid nanostructure of *Parides sesostris* was reported to be templated by hierarchical woven fibrillar structures composed of an as-yet-unidentified material [6].

Despite substantial progress in the structural and optical characterisation of these photonic nanostructures, the molecular composition of butterfly scales remains poorly resolved, limiting our understanding of their assembly and their biomimetic reconstruction. Butterfly cuticle is a composite material comprising chitin, proteins, lipids and pigments [7], yet it is often described simply as “chitinous cuticle” in the absence of detailed molecular information. Variation in the composition and spatial distribution of these constituents influence cuticular organization and, consequently, its optical and functional properties. Proteins and lipids are integral to butterfly scale development and are present within the scale-forming epithelial cells. Cytoskeletal proteins, including F-actin, contribute to the formation and organisation of scale nanostructures. Pioneering developmental transmission electron microscopy studies by Ghiradella [8] further demonstrated that extensive membrane remodelling precedes cuticle maturation [5], implicating lipids as an additional but largely unexplored molecular component of photonic-scale morphogenesis. However, the few lipidomic studies conducted in butterflies have generally reduced intact lipid species to their constituent fatty acids, primarily to investigate migration or the uptake of host-plant-derived lipids [9–12]. This omission is particularly notable because amphiphilic lipids can spontaneously self-organise in aqueous environments into lipidic lyotropic liquid-crystalline phases. Some of these phases form highly curved, bicontinuous cubic structures with three-dimensional periodicity resembling nanostructures found in biological photonic systems, although typically at length scales approximately one order of magnitude smaller [13, 14]. Proteomic analyses of butterfly wing-scale tissues are similarly lacking. Spatially resolved characterisation of intact lipids and proteins could therefore reveal the molecular landscape of structurally coloured scales and connect developmental processes to the nanostructure of the mature photonic material.

Here, we address this gap using *Parides eurimedes mylotes*, a neotropical butterfly with a sex-specific optical phenotype generated by a one-dimensional Bragg-stack photonic crystal. This species belongs to the same family as gyroid-forming butterflies but is more readily available and also forms its photonic nanostructure within the scale lumen, providing an experimentally accessible system in which to investigate the molecular basis of photonic-scale organisation. We combine region-resolved lipidomics and proteomics to identify molecular species in defined wing regions. Whereas previous molecular studies have predominantly analysed whole adult butterflies, our tissue-resolved approach distinguishes structurally coloured regions from pigmented regions and compares their molecular composition between sexes. By integrating these molecular profiles with the nanostructures observed in the corresponding scale types, we resolve the molecular landscape associated with distinct optical phenotypes and link scale composition to structural organisation. Furthermore, we provide a complete protocol for simultaneous lipidic and proteomic analyses in insects.

More broadly, identifying the lipids, proteins and structural components involved in the formation of structurally coloured scales provides insight into the biological mechanisms that organise hierarchical (optical) materials. Such knowledge will enable biomimetic strategies that extend beyond structural colouration, including the bottom-up fabrication of functional, hierarchical three-dimensional nanomaterials. Our integrated framework lays the foundation for single-cell and developmentally dependent, spatially resolved multi-omics to connect molecular composition to material organisation in non-model insects and for translating biological assembly principles into cell-free nanobiotechnology.

## Results

### Region-resolved molecular profiling of pigmentary and structurally colored wing regions

We first characterised the nanostructures underlying the colored patches on the upper wing of *Parides eurimedes mylotes* by transmission electron microscopy (TEM). The male green patch has a multilayer structure within its lumen, whereas the female yellow patch lacks this photonic structure (Figure 1a). We then dissected each specimen into four parts: body, forewing, colored patch, and hindwing (Figure 1b). The tissues were lyophilised and mechanically homogenised for omics analyses.

**Fig. 1.**
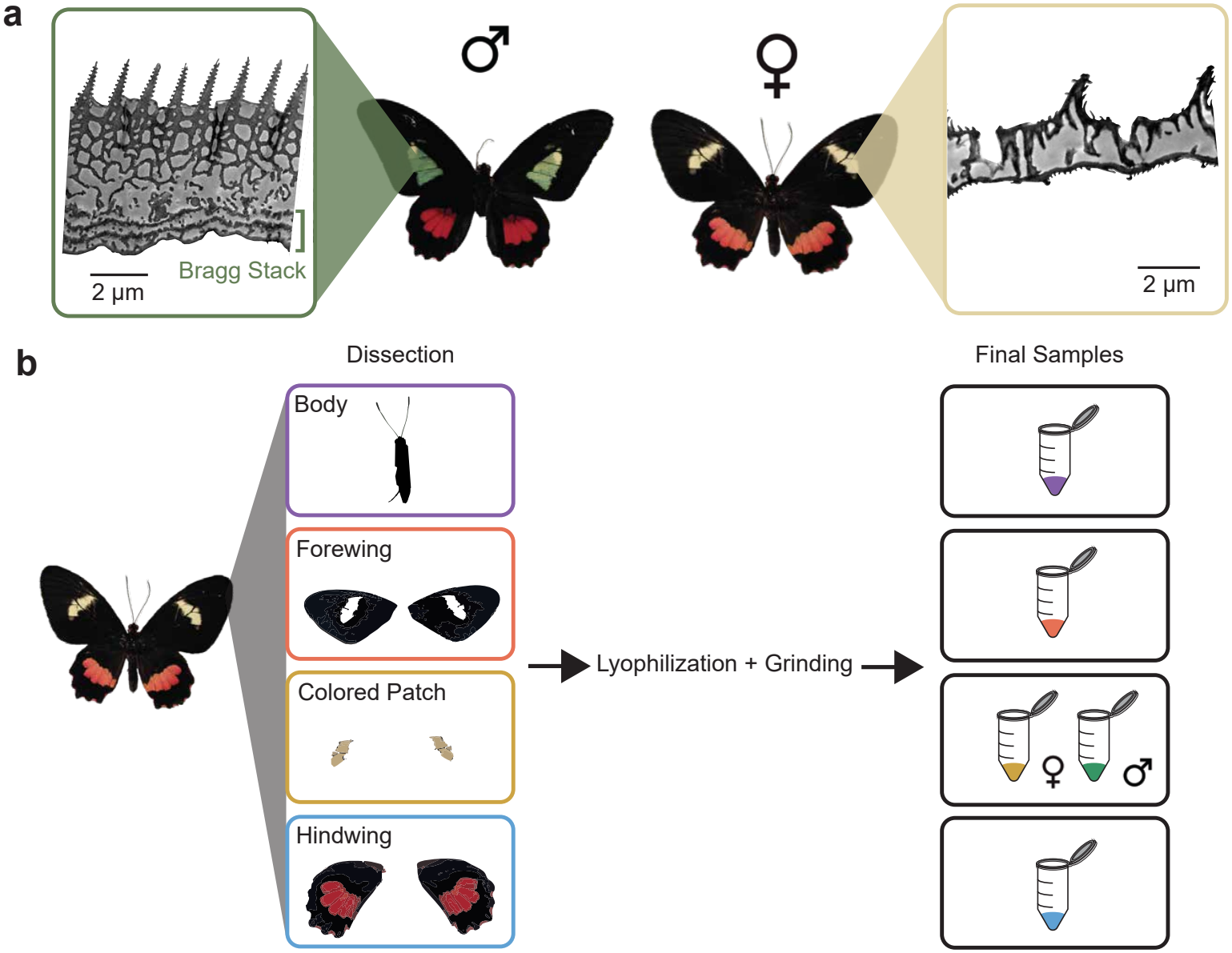
Spatial sampling of butterfly wing tissues and sample preparation for molecular profiling. **a**, Male and female *Parides eurimedes mylotes* butterflies, and transmission electron microscopy images of wing scale cross sections showing the underlying nanostructures of the colored patches in the upper wing. **b**, Spatial dissection of the butterfly into four parts (the body, forewing, colored patch, and the hindwing) and the processing of the corresponding tissue samples for multi-omic analyses.

### Integrated spatial lipidomic and proteomic analyses

The small, spatially restricted samples limit the material available, and splitting each sample between lipidomics and proteomics adds variation. Conventional fatty acid analyses also discard information on lipid class and acyl chain composition. We therefore established an integrated workflow: the homogenised sample was first subjected to solvent-based lipid extraction (Figure 2a), and the remaining tissue pellet was processed for protein extraction (Figure 2b), recovering both lipids and proteins from the same sample. Both extracts were analysed by liquid chromatography–tandem mass spectrometry (Figure 2c). This workflow extracts complementary biochemical information from a spatially heterogeneous sample while preserving intact lipid information, without separate specimens for each omics study.

**Fig. 2.**
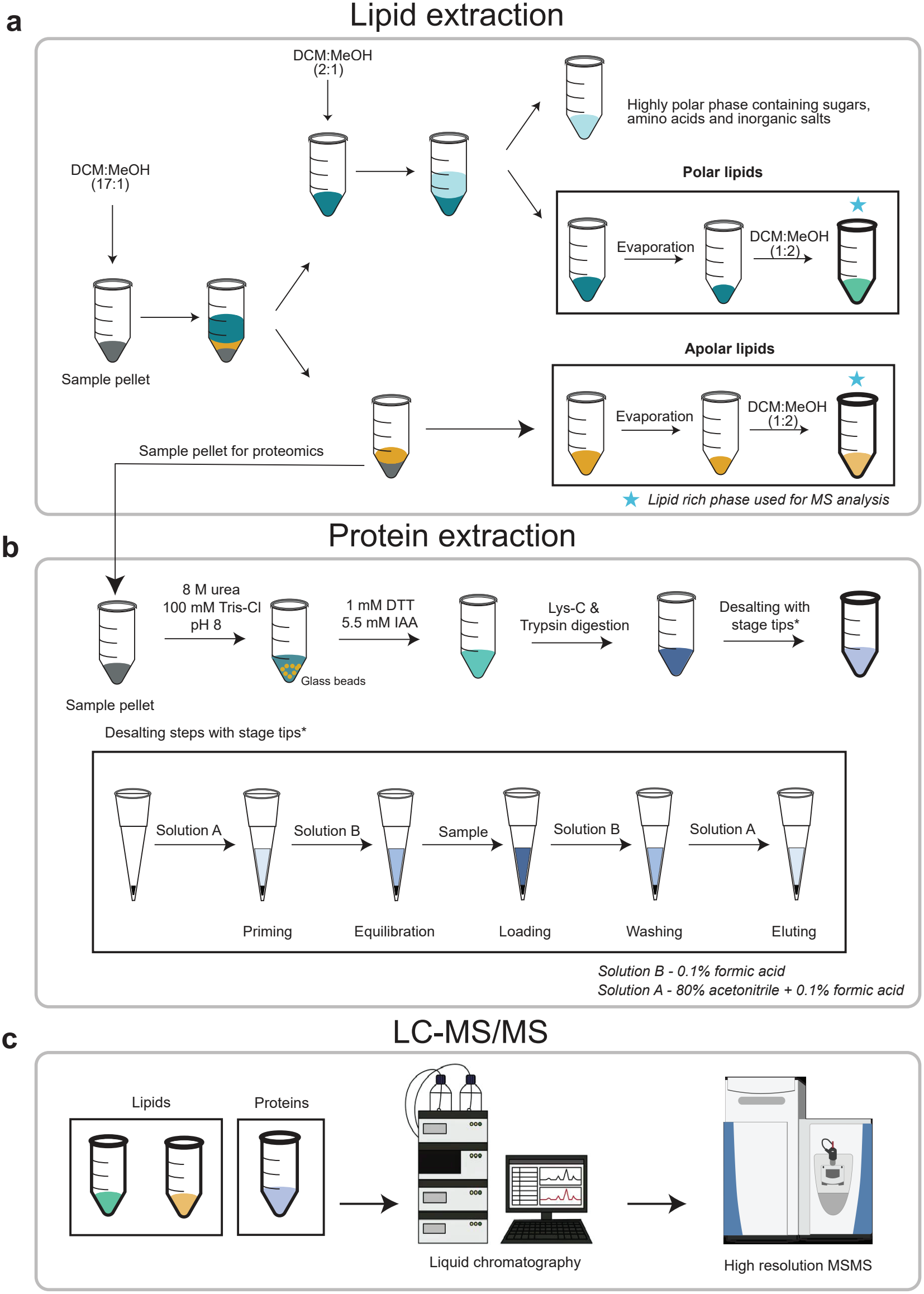
Integrated workflow for spatially resolved lipidomic and proteomic profiling. **a**, Workflow for lipidomic analysis of spatially dissected wing regions. **b**, Workflow for proteomic analysis of corresponding spatially defined tissues **c**, Downstream liquid chromatography-tandem mass spectrometry workflow.

### Region-specific lipidomic signatures distinguish the pigmented and the structurally colored parts

Untargeted lipidomics revealed distinct molecular signatures between the green and yellow patches. Principal component analysis (PCA) captured 47.6 % of the variance (PC1 28.5 %, PC2 19.1 %) and separated the photonic green scales of males from the pigmented yellow scales of females into largely distinct clusters (Figure 3a). Differential abundance analysis (Figure 3c; significance threshold *p* = 0.05) showed that features specific to the green patch were enriched in triacylglycerols (TAG), diacylglycerols (DAG), and wax monoesters, whereas the female pigmented region showed purine alkaloid and halogenated fatty acid features. Halogenated fatty acids also appeared in the blanks and are considered analytical contaminants.

**Fig. 3.**
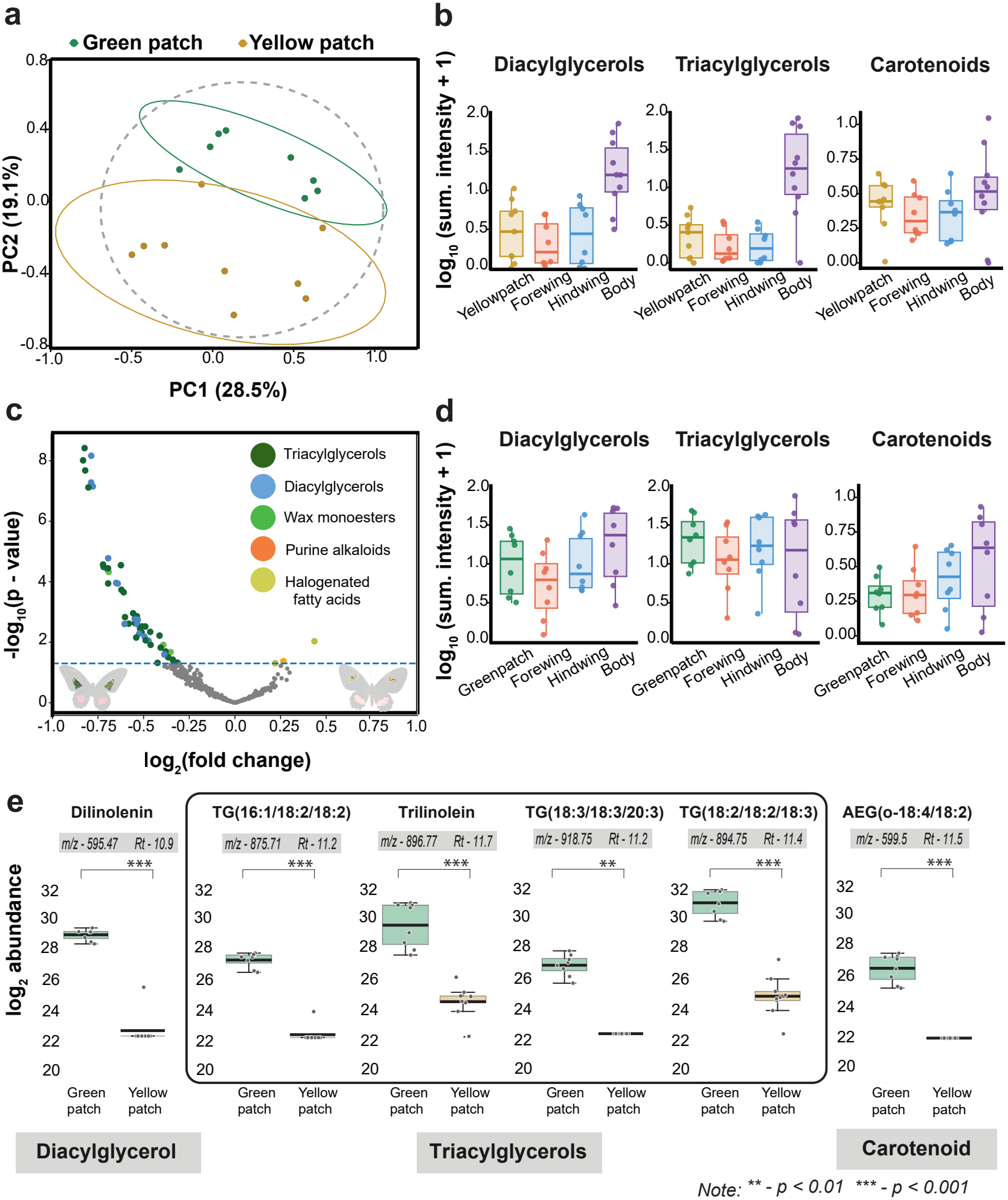
Feature-based molecular network (FBMN) reveals region-specific lipid features in butterfly wings. **a**, PCA of lipidomic profiles from green (photonic) and yellow (pigmented) wing patches. **b**, Relative abundance of diacylglycerols, triacylglycerols, and carotenoids across the body, upper wing, lower wing, and yellow patch in the females. **c**, Differential lipid features between green (on the left) and yellow (on the right) patches. **d**, Relative abundance of diacylglycerols, triacylglycerols, and carotenoids across the body, upper wing, lower wing, and yellow patch in the males. **e**, Candidate lipid features enriched in the green patch. Bar plots showing the six most abundant greenpatch-associated features. For each feature, the annotated name, *m*/*z* ratio, and retention time (*R*_t_) are above the bar, and the assigned lipid class is below the bar.

We compared the relative abundance of DAG, TAG, and carotenoids across the colored patches, forewings, hindwings, and body. In females (Figure 3b), DAG and TAG were more abundant in the body than in the wing tissues (0.49 and 0.46 in the yellow patch), and carotenoids varied less between parts (0.51 in the body, 0.41 in the yellow patch). In males (Figure 3d), DAG was slightly higher in the body than in the wings, TAG was highest in the green patch (1.39), and carotenoids were more abundant in the body and hindwing than in the forewing and green patch.

Among the significant green-patch features (Figure 3e) were the DAG dilinolenin (*m*/*z* = 595.47, retention time 10.9 min) which was detected as a dehydrated protonated ion [*M* + *H− H*_2_*O*]^+^; several TAGs, including TG(16:1/18:2/18:2), trilinolein, TG(18:3/18:3/20:3), TG(18:2/18:2/18:3); and the putative ether-linked glycerolipid AEG(o-18:4/18:2) was detected as the protonated ion [*M* +*H*]^+^. Numbers in parentheses give chain length and number of double bonds, and ‘o’ denotes an ether-linked chain; TG(16:1/18:2/18:2), for example, comprises one 16-carbon monounsaturated and two 18-carbon diunsaturated chains.

### Chemical validation supports the identification of candidate lipid features

#### Compounds specific to pigmented regions

A feature at *m*/*z* = 551.4246 (Wikidata: Q422067) was detected as a dehydrated protonated ion [*M* +*H − H*_2_*O*]^+^ (Figure 4a), present only in the red patches, and was annotated as a carotenoid. Lutein, proposed as the candidate, matched the standard in retention time and MS/MS spectrum, with a cosine score of 0.85. A feature at *m*/*z* = 565.4044 (Wikidata: Q385657) detected as a protonated ion [*M* + *H*]^+^(Figure 4b), present in both yellow and red patches, was also annotated as a carotenoid, and we hypothesized it to be canthaxanthin. The standard and sample showed matching chromatographic peaks at around 5.9 and 6.1 min, with a cosine similarity of 0.94 (where 1 represents identical spectra), confirming the presence of canthaxanthin in the yellow and red patches.

**Fig. 4.**
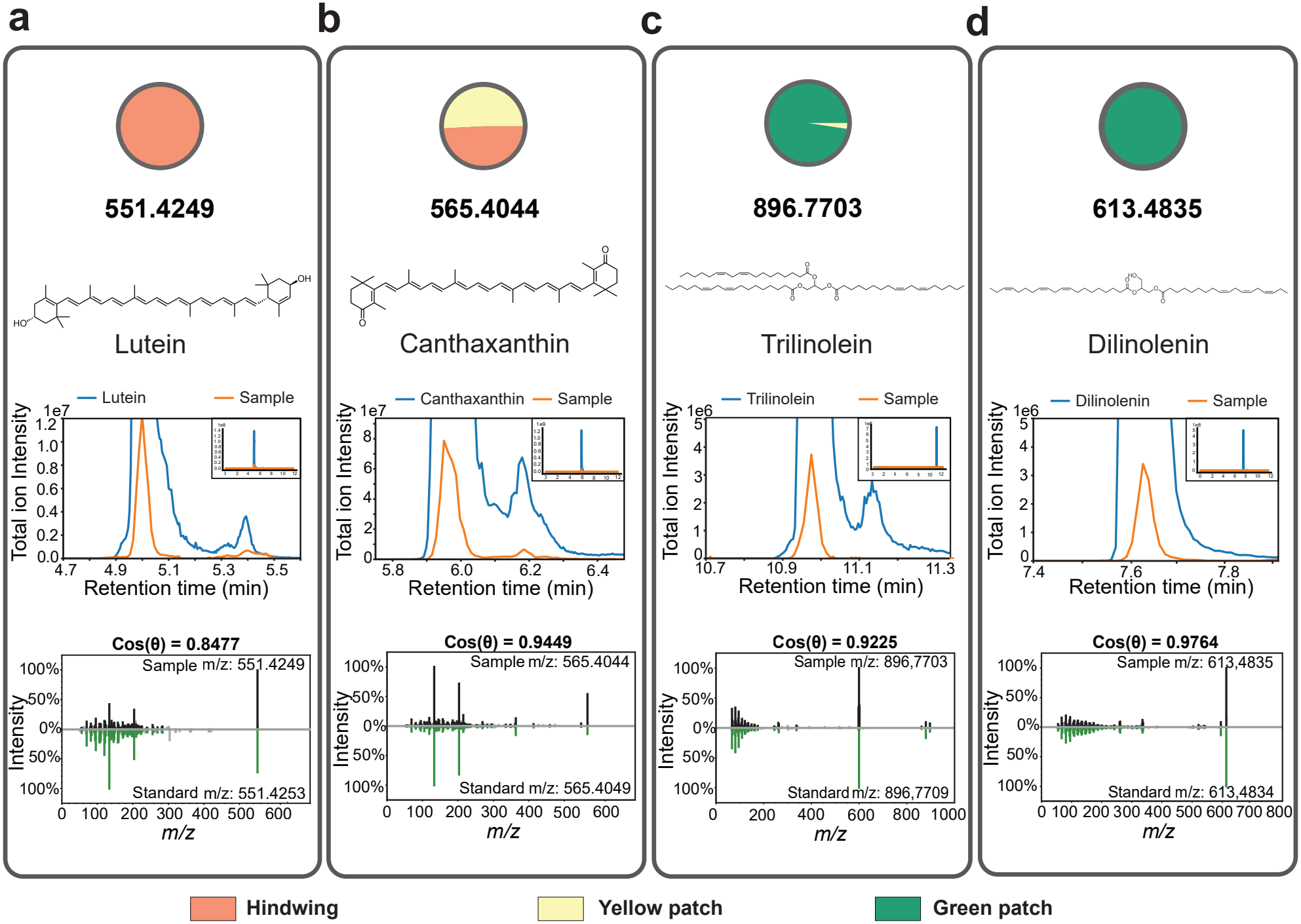
Chemical and optical validation of candidate pigment and lipid annotations. **a**, Node indicating the putative lutein feature, its molecular structure, the LC retention-time matching for lutein standard and sample, and MS/MS spectral comparison between lutein standard and sample. **b**, Node indicating the putative canthaxanthin feature, its molecular structure, the LC retention-time matching for canthaxanthin standard and sample, MS/MS spectral comparison between canthaxanthin standard and sample. **c**, Node indicating the putative trilinolein feature, its molecular structure, the LC retention-time matching for trilinolein standard and sample, MS/MS spectral comparison between trilinolein standard and sample. **d**, Node indicating the putative dilinolenin feature, its molecular structure, the LC retention-time matching for dilinolenin standard and sample, MS/MS spectral comparison between dilinolenin standard and sample.

#### Compounds present in the photonic regions

Many annotated TAG clusters and nodes were present mostly in the green patches. We confirmed the candidate trilinolein by retention-time overlap and by mirror-image overlap of the precursor-ion fragmentation pattern (cosine score: 0.92; Figure 4c), and likewise the DAG dilinolenin (cosine score: 0.97; Figure 4d).

### Region-specific proteomic analysis reveals differential enrichment of cytoskeletal and supramolecular structural proteins

Proteomic analysis of the pellets remaining after lipid extraction identified 6500 proteins across the tissues, with substantial overlap; the body contained the most (Figure 5a). Using the species-specific genome of *Parides eurimedes mylotes* instead of that of *Bombyx mori* increased protein identifications by 50 % [15]. In PCA, two components captured 75 % of the variance (PC1 67.7 %, PC2 7.3 %); body samples formed a distinct cluster along PC1, whereas the wing tissues overlapped substantially (Figure 5b).

**Fig. 5.**
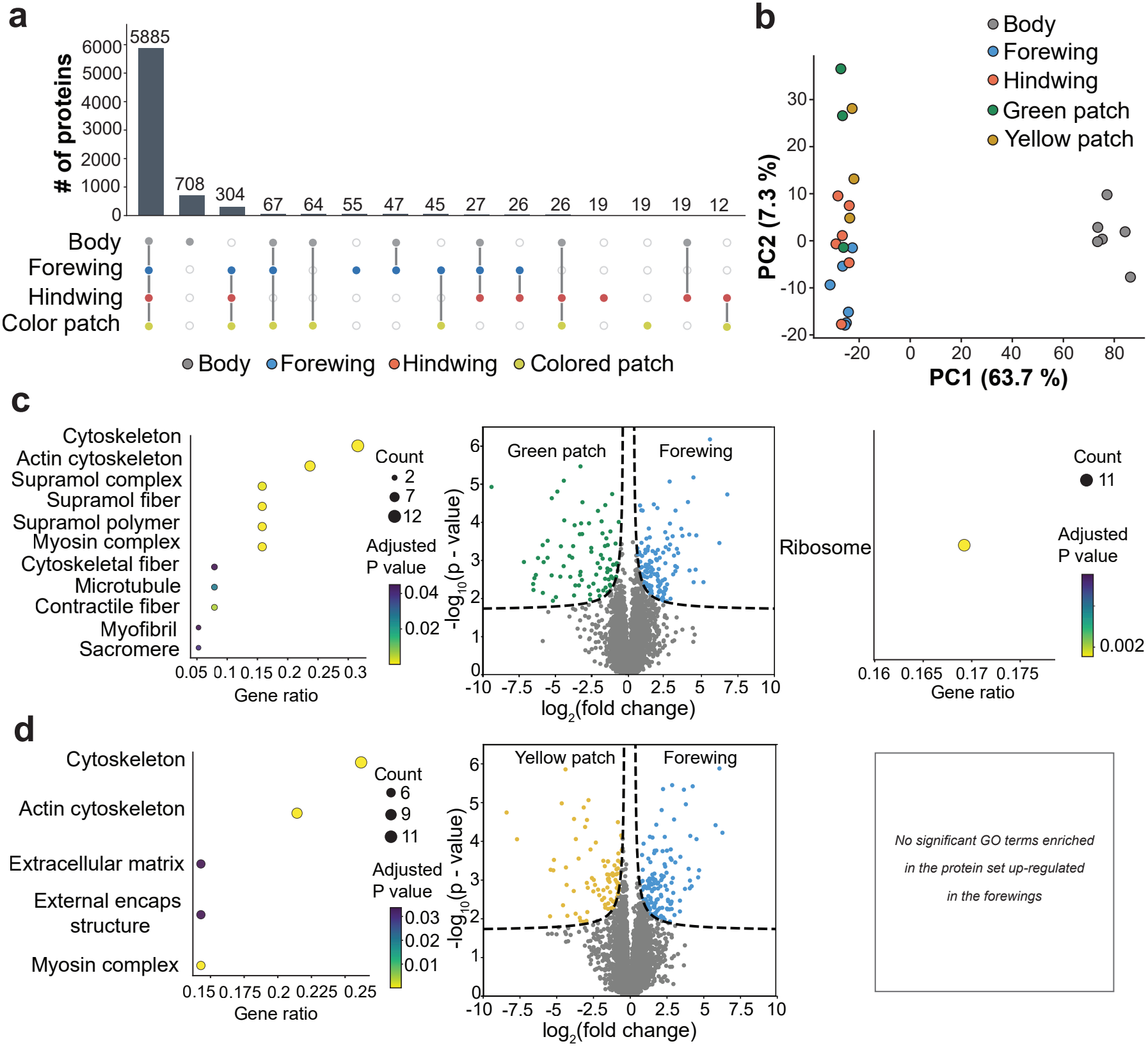
Region-specific proteomic profiling resolves distinct molecular states across the different butterfly body parts. **a**, Upset plot showing the overlap of proteins identified across the body, forewing, hindwing, and colored patch, including both the pigmented and photonic patches. **b**, Principal component analysis of the proteomic profiles, across body, forewing, hindwing, green, and yellow patch. **c**, Differential proteomic analysis of male green patch versus the black forewing, with the corresponding GO-enriched dot plots associated with each region. **d**, Differential proteomic analysis of female yellow patch versus the black forewing, with the corresponding GO-enriched dot plots associated with each region. Volcano plots show a black curved line corresponding to FDR < 0.05, and GO enrichment was filtered at FDR < 0.05 and at least 2-fold enrichment.

We compared the male green patch with the black forewing as background reference (FDR < 0.05, absolute fold change of 2; Figure 5c). Proteins more abundant in the green patch were enriched for the cellular component terms cytoskeleton, actin, myosin complex, supramolecular polymer, supramolecular fibre, supramolecular complex, contractile fibre, microtubule, sarcomere, and myofibril, whereas those abundant in the black forewing were enriched in ribosomal proteins. Enriched biological process and molecular function terms are given in Figures S1 and S2. Under the same thresholds, proteins more abundant in the female yellow patch than in the black forewing were enriched for cytoskeleton, actin, myosin, external encapsulating structure, and extracellular matrix. In contrast, no GO terms were significantly enriched in the black forewing (Figure 5d). Both coloured patches thus show enrichment of cytoskeletal-associated actin and myosin proteins relative to the black forewing, with broader functional differences between the sexes.

### Cross-comparison reveals a conserved proteomic signature in the green patch

Comparing the green patch separately with the yellow patch and the black forewing, 64 and 116 proteins, respectively, were more abundant in the green patch, with six at the intersection between the two comparisons (Figure 6a). Related GO terms were manually consolidated into broader categories (Figure 6b), and the abundance profiles of three functional groups are shown as heatmaps (Figure 6c). Lipid metabolism terms (lipid acyltransferase, O-acyltransferase and lipid kinase activity, prostaglandin metabolism) and vesicle-mediated transport and membrane trafficking terms (receptor-mediated endocytosis, transmembrane transport, SNARE binding, intracellular transport, SNAP receptor system) were more abundant in the male green patch and male forewing. The first two lipid terms were not detected in the female yellow patch. Carbohydrate metabolism terms (UDP-glycosyltransferase activity, UDP-glucose synthesis, carbohydrate transport) were most abundant in the male and female forewings, followed by the green patch, and least in the yellow patch.

**Fig. 6.**
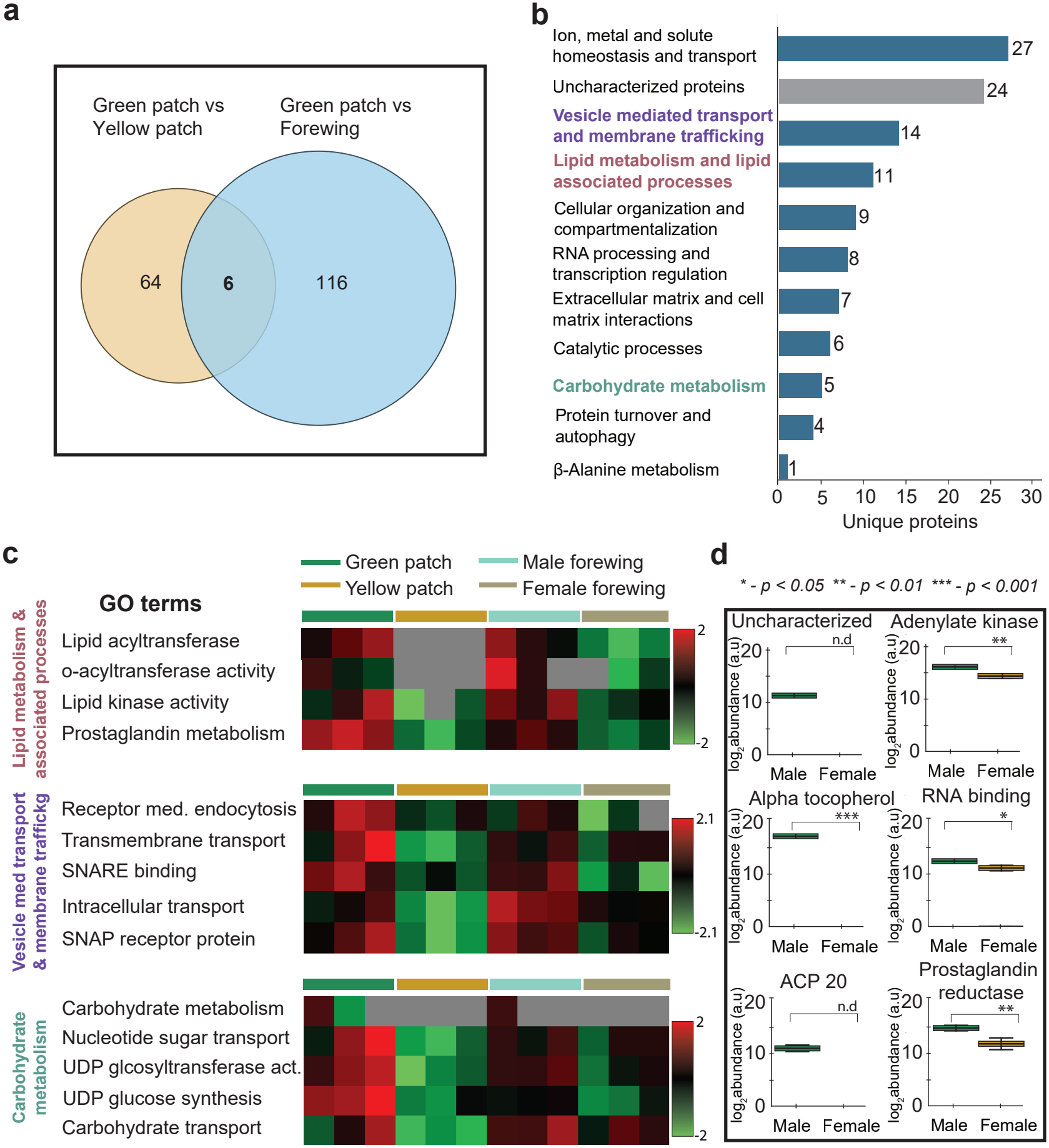
Integrated proteomic signatures associated with the green patch. **a**, Venn diagram showing proteins more abundant in the green patch than in the yellow wing patch (64) and black forewing (116), with six proteins shared between the two comparisons. **b**, Manual curation and consolidation of the abundant GO terms in the male green patch compared to the female yellow patch (70). **c**, Heatmaps showing the abundance of proteins annotated with lipid metabolism and lipid-associated processes (in burgundy), vesicle-mediated transport and membrane trafficking (in purple), and carbohydrate metabolism (in blue). **d**, Relative abundance of six proteins shared between the two green patch comparisons across male green and female yellow patches. Statistical significance was assessed by pairwise comparisons, with *p* < 0.05, *p* < 0.01, and *p* < 0.001 indicated by one, two, and three asterisks, respectively.

The six shared proteins, all more abundant in the green than in the yellow patch (Figure 6d), were an uncharacterized protein, adenylate kinase, alpha-tocopherol protein-like, an RNA-binding protein, adult-specific cuticular protein (ACP 20), and prostaglandin reductase (sequences in Supplementary Data 1. Protein sequences). Two of these have lipid-associated functions, linking the proteomic and lipidomic datasets.

## Discussion

Colouration in butterfly wings arises from molecular composition, hierarchical organisation, and pigment localisation, which are rarely resolved within the same tissue. Combining spatially resolved lipidomics and proteomics, we found pronounced molecular differences between the green photonic patch and the yellow pigmentary patch.

Lepidopteran scales arise from regionally specified precursor cells [16], and their development coordinates cytoskeletal organisation, cuticle deposition, and sclerotisation. Lipids are established cuticle components transported from epidermal tissues [17–19], so distinct lipidomic signatures across wing tissues are consistent with differential lipid incorporation during development. The photonic colored patch was found to be rich in DAG and TAG. In contrast, the pigmentary patch had relatively higher carotenoid levels, indicating that the sex-specific coloured regions occupy distinct biochemical states. Previous lipidomic studies of butterflies and caterpillars examined constituent fatty acids rather than intact lipids. Our workflow preserves acyl chain length and degree of unsaturation, which matters here because long-chain DAG and TAG are prominent in the photonic crystal region. Such intact neutral lipids are non-volatile and thus unlikely to act directly as volatile pheromones.

Matching to authentic standards confirmed the presence of canthaxanthin and lutein in the yellow and red scales. This distinguishes *Parides eurimedes mylotes* from *Papilio* species, where papiliochromes colour yellow and reddish-brown scales [20], and from nymphalids, where ommochromes and 3-hydroxykynurenine are associated with yellow and red colouration [21]: in *Parides*, carotenoids account for both.

The proteomic data adds a complementary dimension. The pigmented region was enriched in cytoskeletal, actin, myosin, external encapsulating structure, and extracellular matrix proteins, and the photonic patch additionally showed broader enrichment of cytoskeletal, contractile, and supramolecular fibres, complexes, and polymers, microtubules, sarcomeres, and myofibrils. This is consistent with studies showing that the actin cytoskeleton is key to nanostructure development [22]. Disrupting actin polymerisation in *Heliconius sara* abolished photonic properties but did not resolve which proteins participate [23]. By identifying nine actin proteins enriched in the photonic patch, with high sequence confidence from species-specific genomic data, we move from pathway-level perturbation to molecular composition, providing candidates for targeted study. Relative to the yellow patch, the green patch also showed more GO terms for lipid metabolism, membrane trafficking, carbohydrate metabolism, and cuticular components, suggesting a role for these processes in the development of the male photonic region.

These findings support the hypothesis that structural colour arises from the spatial organisation of specific molecules from designated pathways and their hierarchical assembly. The enrichment of TAG and DAG in the photonic patch suggests that regional lipid composition may influence membrane organisation during nanostructure development: DAGs modulate membrane curvature and promote inverse cubic phases in lipid–water systems [24], whereas the role of TAGs is less established. However, our measurements were performed on mature scales and do not establish a causal role of individual lipid species in gyroid formation. Targeted validation and spatial localisation of candidate proteins at the single-cell level and over development, together with functional perturbation of candidate pathways, will be needed to link molecular organisation to the final photonic structure.

In summary, we established an integrated region resolved lipidomics–proteomics workflow for insects that minimises the material required across defined butterfly wing regions. It revealed distinct TAG and DAG clusters in the photonic scales, identified the carotenoids responsible for red and yellow colouration, and showed extensive differences in cellular components (supramolecular fibres, complexes, and polymers, cytoskeletal fibres, microtubules, contractile fibres, and sarcomeres) between the photonic region and the black forewing. The photonic region also showed enrichment of proteins associated with lipid metabolism, vesicle-mediated transport, and membrane trafficking, consistent with the female yellow patch and reflecting regional lipid differences. Butterfly wing colouration is thus accompanied by coordinated spatial differentiation in lipid composition, pigment chemistry, and structural proteins. This work establishes the molecular library from which more specific molecular targeted studies can follow to understand the formation of diverse functional nanostructures ultimately leading to their biomimetic recreation.

## Methods

### Sample processing

Individuals were separated into four fractions: upper wings, lower wings, coloured patches of the upper wings (green in males and yellow in females), and the body. Samples were freeze-dried for 3 days to eliminate residual moisture (Figure 1). Dried tissues were ground into a fine powder using corrugated steel beads in a mechanical shaker for a short time. Approximately 30 mg of powdered material was transferred into a 2 mL Eppendorf tube for extraction.

### Lipidomic analysis

#### Lipidomic extraction

A modified Schevchenko protocol to extract the lipids was performed on dissected butterfly samples. Approximately 30 mg of powdered material was transferred into a 2 mL Eppendorf tube for extraction. A solvent mixture of 495 µL dichloromethane:methanol (17:1, v/v) and 100 µL water was added to the powdered sample. The mixture was vortexed for 15 s, sonicated for 5 min, and centrifuged at 16 000 rev min^*−*1^ for 5 min. The upper phase was manually transferred to a separate vial using a pipette, and the lower phase was dried under a gentle stream of N_2_ gas.

The upper phase was again re-extracted with 495 µL dichloromethane:methanol (2:1, v/v) without the addition of water. The mixture was centrifuged at 16 000 rev min^*−*1^ for 5 min. The upper phase was transferred to a GC vial, and the lower phase was dried under N_2_ gas [25, 26].

The dried residues from both extraction steps were resuspended in 100 µL dichloromethane:methanol (2:1, v/v) and transferred to GC vials for subsequent GC–MS analysis. A blank was processed under the same conditions to detect any potential contamination during the extraction steps.

### Data Aquisition

#### LC-MS parameters

Chromatographic separation was performed on a Vanquish Flex UPLC system (Thermo Fisher Scientific) interfaced with a Q-Exactive Plus mass spectrometer (Thermo Fisher Scientific), using a heated electrospray ionisation (HESI-II) source. Thermo Scientific Xcalibur 3.1 software was used for instrument control. The LC conditions were as follows: column, Waters BEH C18 100 mm × 2 mm, 1.7 µm; mobile phase, (A) water with 0.1 % formic acid; (B) acetonitrile with 0.1 % formic acid; flow rate, 600 µL min^−1^; injection volume, 2 µL; gradient, linear gradient of 2 % to 100 % B over 10 min and isocratic at 100 % B for 2 min. The optimised HESI-II parameters were as follows: source voltage, 3.5 kV (pos); sheath gas flow rate (N_2_), 55 units; auxiliary gas flow rate, 15 units; spare gas flow rate, 3.0; capillary temperature, 350 ^*?*^C; S-Lens RF Level, 50. The mass analyser was calibrated using a mixture of caffeine, methionine-arginine-phenylalanine-alanineacetate, sodium dodecyl sulfate, sodium taurocholate, and Ultramark 1621 in an acetonitrile/methanol/water solution containing 1 % formic acid by direct injection. Data-dependent tandem mass spectrometry (MS/MS) events were acquired on the three most intense ions detected in the full-scan MS (Top3 experiment). The MS/MS isolation window width was 1 Da, and the stepped normalised collision energy was set to 15, 30, and 45 units. In data-dependent MS/MS experiments, full scans were acquired at a resolution of 35 000 FWHM (at *m*/*z* = 200) and MS/MS scans at 17 500 FWHM, both with an automatically determined maximum injection time. After being acquired in an MS/MS scan, parent ions were placed in a dynamic exclusion list for 3 s.

### Data processing

The MS data were converted from “.RAW” (Thermo) standard data format to “.mzML” format using the MSConvert software, part of the ProteoWizard package [27]. The converted files were processed using the MZmine software suite v. 2.53 [28]. The parameters were adjusted as follows: the centroid mass detector was used for mass detection, with the noise level set to 2.0 × 10^4^ for the MS level set to 1, and to 0 for the MS level set to 2. The ADAP chromatogram builder [29] was used and set to a minimum group size of 5 scans, a minimum group intensity threshold of 2.0 × 10^4^, a minimum highest intensity of 2.0 × 10^4^, and an *m*/*z* tolerance of 12 ppm. For chromatogram deconvolution, the algorithm used was wavelets (ADAP). The intensity window signal-to-noise (*S*/*N*) was used as an S/N estimator with an S/N ratio set at 15, a minimum feature height at 2.0 × 10^4^, a coefficient area threshold at 80, a peak duration range from 0.02 min to 1.00 min and the retention time (RT) wavelet range from 0.02 min to 0.05 min. Corresponding MS2 was paired with the following parameters (0.025 Da and 0.15 min). Isotopes were detected using the isotope peaks grouper with an *m*/*z* tolerance of 8 ppm, an RT tolerance of 0.08 min (absolute), the maximum charge set at 4, and the lowest *m*/*z* was used as the representative isotope. Peak alignment was performed using the join aligner method (*m*/*z* tolerance at 12 ppm), absolute RT tolerance 0.08 min, weight for *m*/*z* at 30, and weight for RT at 30.

### Molecular networking

To analyse the spectral diversity of the profile collection, a molecular network (MN) was created on the GNPS website (http://gnps.ucsd.edu) using the “.mgf” spectra file generated at the previous step [30]. The precursor ion mass tolerance was set to (0.02 Da and an MS/MS fragment ion tolerance of (0.02 Da. A network was then created where edges were filtered to have a cosine score above 0.7 and more than six matched peaks. Further, edges between two nodes were retained in the network if and only if each node appeared in the other’s top 10 most similar nodes. Finally, the maximum size of a spectral family was set to 100, and the lowest-scoring edges were removed from molecular families until the molecular family size was below this threshold. The spectra in the network were then searched against GNPS spectral libraries. All matches kept between network spectra and library spectra were required to have a score above 0.7 and at least six matched peaks. The resulting MN is available online at https://gnps2.org/result?task=4ac40c44175c482da185bf70df88366d&viewname=librarymatches&resultdisplay_type=task. The converted mass spectrometry data files, the corresponding metadata table, and the metabolite annotation results table, together with a Cytoscape file representing the full MN annotated with experimental and theoretical spectral matches, are available under the MassIVE ID https://massive.ucsd.edu/ProteoSAFe/dataset.jsp?accession=MSV000104073.

### Metabolite Annotation

Experimental spectral libraries search: The full spectral data set corresponding to the extracts collection has been uploaded to the MassIVE repository https://massive.ucsd.edu/ProteoSAFe/dataset.jsp?accession=MSV000104073, and a continuous identification workflow is automatically carried out against the Global Natural Products Social Molecular Networking (GNPS) experimental spectral libraries: https://gnps2.org/result?task=4ac40c44175c482da185bf70df88366d&viewname=librarymatches&resultdisplay_type=task.

Theoretical spectral libraries search: In addition to experimental spectral libraries search, we have shown that spectral matching against theoretical spectral libraries of natural products was an efficient way to cover a much wider, yet relevant, spectral space [31]. Furthermore, we showed that taking into account the taxonomical distance between the biological source of the candidate structure and the biological source of the annotated extracts greatly improved the overall quality of the annotation results [32]. Thus, in addition to the spectral search performed at the molecular networking step against publicly available spectral libraries (see previous section) a taxonomically informed metabolite annotation was performed. For this, we first established a large theoretical spectral database of natural products following a previously established metabolite annotation workflow. This spectral database and the associated biological sources metadata were constructed using chemical structures and information compiled during the LOTUS Initiative’s first project, which aimed to establish an open and evolving resource compiling natural products and biological occurrences [33, 34]. The theoretical spectral database is publicly available [35]. The biological sources metadata are available online [33, 34]. The taxonomically informed metabolite annotation was performed using the met_annot_enhancer scripts version v0.1: https://github.com/mandelbrot-project/met_annot_enhancer/releases/tag/v0.1. The parameters used in the taxonomically informed metabolite annotation process, along with the resulting tables, are available at https://gnps2.org/result?task=4ac40c44175c482da185bf70df88366d&viewname=librarymatches&resultdisplay_type=task. Sirius [36] (v.5.5.7) and CANOPUS [37–39] were also employed to proceed to metabolite annotation and attribution of chemical classes to MSMS spectra.

### LC–MS/MS matching to authentic standards

MZmine version 4.10.6 was used for mass extraction based on the molecular formulas of the standards and quality control (QC) samples from the experiment. Extracted chromatograms were generated for visualisation and preparation of the corresponding figures. For the MS comparison, the GNPS2 dashboard (Metabolomics USI: https://metabolomics-usi.gnps2.org/dashinterface/?usi=) was used. The feature numbers generated during the MZmine processing and included in the resulting feature table were entered into the GNPS2 dashboard for subsequent comparison.

## Proteomic analysis

### Proteomic extraction

Proteins were extracted in 8 M urea, 100 mM Tris-Cl, pH = 8, by sonication, followed by using a bead beater. After centrifugation, proteins were reduced with 1 mM DTT, alkylated with 5.5 mM IAA, and digested by Lys-C for (4 h at room temperature. Before overnight trypsin digestion, the urea concentration was reduced to 100 mM. The next day, the samples were acidified using 50 % TFA (final concentration app. 0.5 %, pH < 2) and peptides were desalted on StageTips [40].

### Data acquisition and processing

LC-MS/MS analyses were performed on an Astral Orbitrap mass spectrometer coupled to a Vanquish NEO UHPLC system (all Thermo Scientific). Peptides were separated on a fused silica HPLC column tip (I.D. 75 µm, New Objective, self-packed with ReproSil-Pur 120 C18-AQ, 1.9 µm (Dr. Maisch) to a length of 25 cm) using a gradient of A (0.1 % formic acid in water) and B (0.1 % formic acid in 80 % acetonitrile in water). The samples were loaded with 0 % B with a flow rate of 500 nL min^*−*1^; the peptides were separated by 5 % – 55 % B within 39 min with a flow rate of 500 nL min^*−*1^. The spray voltage was set to 2.0 kV and the ion-transfer tube temperature was set to 280 ^*?*^C; no sheath or auxiliary gas was used. The mass spectrometer was operated in data-independent mode; after each MS scan (*m*/*z* = 380 980; resolution: 240 000, maximum injection time of 3 ms), 149 MS/MS scans with an isolation width of 4 *m*/*z* with no overlap were performed, covering a range from *m*/*z* = 380 to 980. The normalised AGC target value was set to 300 %, and the HCD collision energy to 25 %. The MS raw files were analysed using Spectronaut version 20 and the directDIA+ workflow, with *Parides eurimedes mylotes* (both haplotypes) and common contaminants, such as keratins, as references. Carbamidomethylcysteine was set as a fixed modification, and protein N-terminal acetylation and methionine oxidation were set as variable modifications. Peptide, site and protein FDR were set to 0.01; minimum peptide length was set to 7, and a maximum of three missed cleavages were allowed using trypsin/P as enzyme specificity.

### GO enrichment analysis

Exported normalised protein abundances of each sample were further analysed using Perseus software [41]. GO-term enrichment analyses of significantly different proteins (t-test, *p*-value < 0.05, at least 2-fold difference) were performed with the clusterProfiler R package.

## Acknowledgements

The authors acknowledge the funding provided to Praveen Kumar Jaya Balaji through the industrial PhD program “Talents for Biobased Economies (T4BBI),” a Marie Sklodowska-Curie COFUND project under the EU Horizon 2020 Research and Innovation (grant number 101034323). BiOrbic, the Bioeconomy SFI Research Center of Ireland, man-ages this program. This research was also supported by the SNSF Ambizione fellowship 233007 awarded to Dr. Viola Vogler-Neuling, the NCCR Bioinspired Materials of the Swiss National Science Foundation (grant number 205603), and the Adolphe Merkle Foundation.

## Ethics declarations

Butterfly pupae were purchased from The Entomologist Ltd., located at Middleton Common Farm in East Sussex. The pupae were purchased as *Parides arcas*. The company is a member of the International Association of Butterfly Exhibitors and Suppliers (IABES), which protects wild butterflies and their habitats by promoting sustainable exhibits and supporting conservation, research, and public education. Carmen Rojas Ugalde, from the Universidad de Costa Rica, analyzed the pupae and identified them as textit Parides eurimedes mylotes (H. W. Bates, 1861). The pupae were raised in a humidity- and temperature-controlled terrarium until they eclosed. They were euthanized by putting them in a paper envelope and freezing them at -20 °C for three days before omics analyses.

### Data availability

The full spectral data set corresponding to the lipid extracts collection has been uploaded to the MassIVE repository https://massive.ucsd.edu/ProteoSAFe/dataset.jsp?accession=MSV000104073. Taxonomically informed metabolite annotation was performed using met_annot_enhancer (v0.1). The scripts are available on GitHub https://github.com/mandelbrot-project/met_annot_enhancer/releases/tag/v0.1, and the annotation parameters and resulting tables are available through GNPS2 https://gnps2.org/result?task=4ac40c44175c482da185bf70df88366d&viewname=librarymatches&resultdisplay_type=task. For the MS comparison, the GNPS2 dashboard (Metabolomics USI: https://metabolomics-usi.gnps2.org/dashinterface/?usi=) was used. The feature numbers generated during the MZmine processing and included in the resulting feature table were entered into the GNPS2 dashboard. The proteomic data is available via ProteomeXchange with identifier PXD087722. The lipidomics and proteomics datasets, together with the corresponding analysis codes, are publicly available on Zenodo under the following DOIs: lipidomics, 10.5281/zenodo.20407699; and proteomics, 10.5281/zenodo.20407699.

### Author contributions

Conceptualization: P.J.-K. and V.V.-N.; Data curation: P.J.-K., C.V.,P.-M.A. and M.Stumpe; Formal Analysis: P.J.-K., P.-M.A. and M.Stumpe; Funding Acquisition: P.J.-K., U.S. And V.V.-N., Investigation: P.J.-K., C.V., M.Stumpe, M.Saba and J.L.; Project administration: V.V.-N.; Resources: P.J.-K. and M.Stumpe; Supervision: U.S. and V.V.-N.; Visualization: P.J.-K. and V.V.-N.; Writing-originial draft: P.J.-K.; Writing-review & editing: P.J.-K., P.-M.A., M.Stumpe, U.S. and V.V.-N.

