## Supplementary Information for "Lipid and supramolecular structural protein signatures distinguish structurally colored from pigmented wing regions in the butterfly *Parides eurimedes mylotes*"

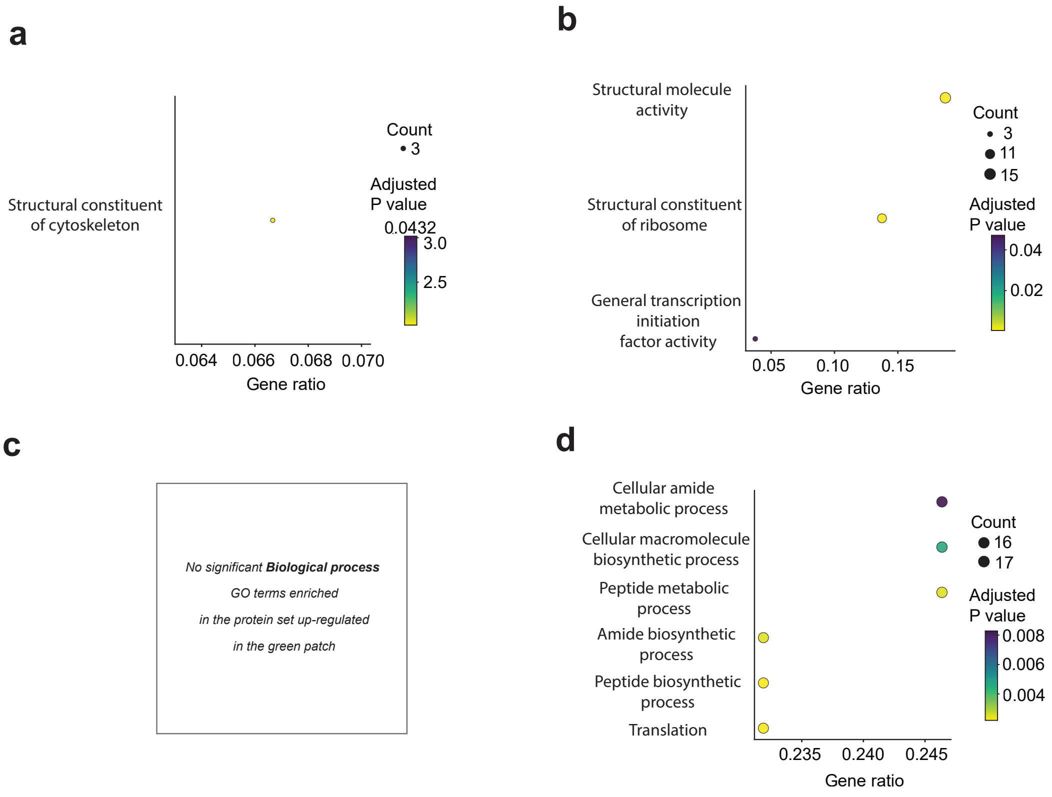


**Fig. 1** Gene Ontology enrichment analysis of male green patch and black forewing proteins. GO enrichment was filtered at FDR < 0.05 and at least 2-fold enrichment. **a**, Molecular function (MF) GO terms enriched in the male green patch. **b**, MF GO terms enriched in the male black forewing. **c**, Biological process (BP) GO terms enriched in the male green patch. **d**, BP GO terms enriched in the male black forewing.

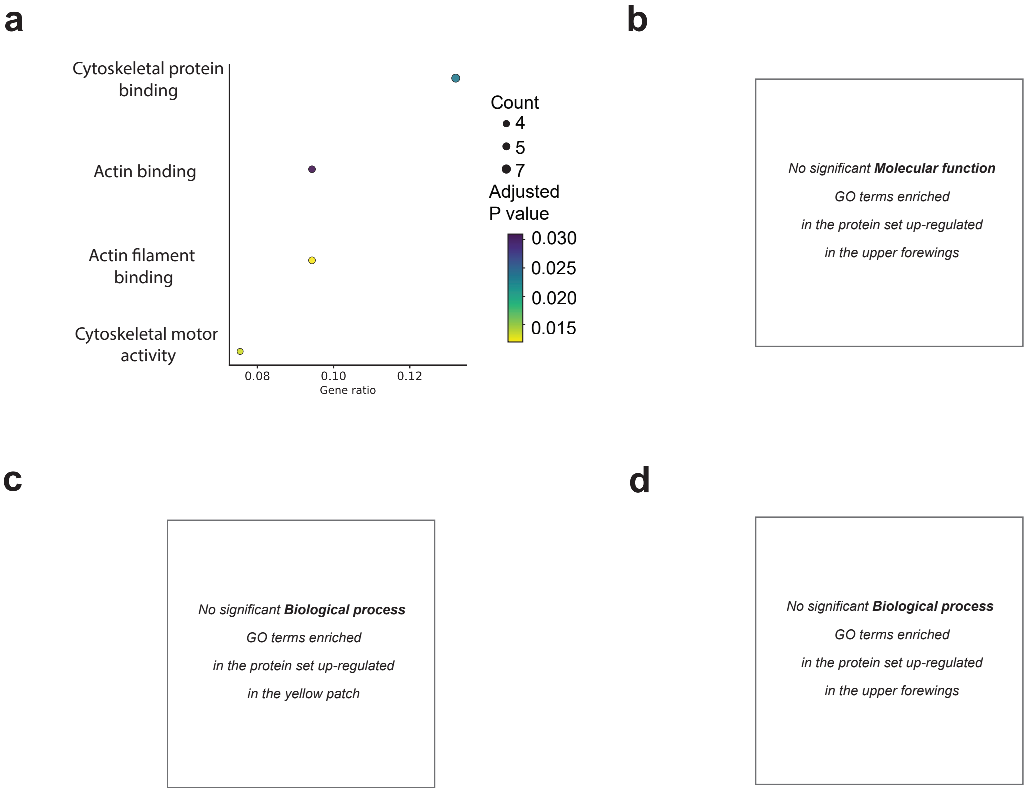


**Fig. 2** Gene Ontology enrichment analysis of female yellow patch and black forewing proteins. GO enrichment was filtered at FDR < 0.05 and at least 2-fold enrichment. **a**, Molecular function (MF) GO terms enriched in the female yellow patch. **b**, MF GO terms enriched in the female black forewing. **c**, Biological process (BP) GO terms enriched in the female yellow patch. **d**, BP GO terms enriched in the female black forewing.

### Supplementary Data 1. Protein sequences

The amino acid sequences of the selected proteins identified in this study are provided below in FASTA format.

```
>gnl|WGS|Hap1_Pmyltmp_007415-P1 uncharacterized protein Pmyltmp_007415
MQTVDEWKVALSVVAEPYTVPDHPRWFGDDLGSVAIYWRGEEGTPPCTLLSRGQGFVAVKWGPLAIIGCY
VSPSKSLADYEEYLDRVAERVTDCLPCPLIVLGDFNAHHRAWGNPKDDRKGDTVLEWAAGLDLRLVNRGS
VSTCVRWQGESVVDLTWATPAAMRMLSGWRVAEEVVTLSDHRHIIFDVAIRKLVADSGRRDGSTTRRWCL
KRLDRDRVEAAACVADWPNAEEVLSDPEEGANWFRETLTQICDASMPRTGRPKRGAVYWWSTEIAQLRKV
CLRSRRQYTRARRKKRRAPNEEIARLYSEYRAATKALQKAVASAKTRSWNELLQGLDRDPWGRPYRIVLG
KLRPWMPPTSEILEPVFLEGVVDTLFPADRLNPSARSHVLSWEWTDETGVTAQELDRAVSRLRARNTAPG
PDGIPGRVWVLSLATLGNRLVELFTSCLRRASFPKPWKEAQLVLLKKEGRAAESPSAYRSICLLDECGKL
FERILAARISRHLSRVGPDLADSQFGFRQGRSTVDAIFRVRSLSEQAVSPGGVALAISLDIVNAFNALPW
DAIREGLVYHDVPPCLRGVIADYLRDRYIVYMDQDGRRIRREIQRGVPQGSVLGPLLWNLAYDAVLRVSL
PAGVNVVCYADDTLVVAGGDNFERAKELAELGTEIVVAKIRELGLEIAPNKTEALWFHKLPRSREPPDSR
VRVGEHEIRVGRYMKYLGLTLDSRWGFEEHFDRLVPRIERIAGAMHRILPNLGGPREKSSQLDILAKELE
HLSAVAPQEKLRDRPDDDDDDDNNNNDDN


>gnl|WGS|Hap1_Pmyltmp_005312-P1 adenylate kinase 1
MDQEATCQPCDEPVVDRKPPCPLQCMSPKCPSYREKPPIDTSKKPIVWVLGGPGSGKGTQCEKIIAKYGF
THLSTGDLLRAEVKSGSDRAKCLTSIMERGELVPNETVLELLKEAIMARAADSKGFLIDGYPREKSQGVA
FENTIAPVTVILYFEASCETLTQRLLGRAASSGRADDNEETIKLRLKTFLDNNDQVLAQYPNKLQRINAE
KSVDEIFGEVQRILDPVVQAAAAN


>gnl|WGS|Hap2_Pmyltmp_007539-P1 adenylate kinase 1
MDQEATCQPCDEPVVDRKPPCPLQCMSPKCPSYREKPPIDTSKKPIVWVLGGPGSGKGTQCEKIIAKYGF
THLSTGDLLRAEVKSGSDRAKCLTSIMERGELVPNETVLELLKEAIMARAADSKGFLIDGYPREKSQGVA
FENTIAPVTVILYFEASCETLTQRLLGRAASSGRADDNEETIKLRLKTFLDNNDQVLAQYPNKLQRINAE
KSVDEIFGEVQRILDPVVQAAAAN


>gnl|WGS|Hap1_Pmyltmp_005461-P1 alpha-tocopherol transfer protein-like
MASTATLVQPSGEMWKRIRVELNEDVNTRDQYLTAIKEWLRKEPHLPDEWDEDRMMTFLRGCKFSLEKCK
RKLDMYFTMRTACPEFFTNRDIKKPELVDLMNRVQGPPMPGLTPNGRRVTVVRGIDKNIDTNHLCDVFKV
```

```
AMMIGDVRLKEEVEGVGGDIYILDASVVSPSHLTKLSPSTIKKFLICVQEAYPVKLKEVHVVNASPIVDK
VVSIVKPFLKEKIKNRIFTHTDFNTLYEYIPKEVLPVEYGGTCGTLKEINDAWVKKLEDYTEWFKEQEEI
KANEALRPGKPTNYDELFGIDGSFRQLAID


>gnl|WGS|Hap2_Pmyltmp_005273-P1 alpha-tocopherol transfer protein-like
MASTATLVQPSGEMWKKIRVELNEDVNTRDQYLTAIKEWLRKEPHLPDEWDEDRMMTFLRGCKFSLEKCK
RKLDMYFTMRTACPEFFTNRDIKKPELVDLMNRVQGPPMPGLTPNGRRVTVVRGIDKNIDTNHLCDVFKV
AMMIGDVRLKEEVEGVGGDIYILDASVVSPSHLTKLSPSTIKKFLICVQEAYPVKLKEVHVVNASPIVDK
VVSIVKPFLKEKIKNRIFTHTDFNTLYEYIPKEVLPVEYGGTCGTLKEINDAWVKKLEDYTEWFKEQEEI
KANEALRPGKPTNYDELFGIDGSFRQLAID


>gnl|WGS|Hap1_Pmyltmp_008598-P1 RNA-binding protein 6 isoform 1-T1
METQSQDVAHSPAEVPNDPGKMFVGGLSWQTSPESLRDYFSKFGEITEVMVMKDPTTRRSRGFGFITFGD
PASVDKVLAQGTHELDGKKIDPKVAFPRRAHPKMVTRTKKIFVGGLSAPTTLEDVKNYFEQFGPIEDAML
MFDKQTNRHRGFGFVTFQSEDIVDKVCEIHFHEINNKMVECKKAQPKEVMLPANLAKTRAAGRGAYDFMW
SLGALPDGFPAAYAAYAAGRSYSGYPSFGLPYPAVDLTNGQLTPNNNTPLLAQALSVMNNYQAAAAGFGP
PASPHAAGGRAGFPAASSPGPALDVYGSAADSVGYVQAASPQPSGFPAIAVSRAPLNYTPGPLIPAAFTN
GYH


>gnl|WGS|Hap2_Pmyltmp_008708-P1 RNA-binding protein 6 isoform 1-T1
METQSQDVAHSPAEVPNDPGKMFVGGLSWQTSPESLRDYFSKFGEITEVMVMKDPTTRRSRGFGFITFGD
PASVDKVLAQGTHELDGKKIDPKVAFPRRAHPKMVTRTKKIFVGGLSAPTTLEDVKNYFEQFGPIEDAML
MFDKQTNRHRGFGFVTFQSEDIVDKVCEIHFHEINNKMVECKKAQPKEVMLPANLAKTRAAGRGAYDFMW
SLGALPDGFPAAYAAYAAGRSYSGYPSFGLPYPAVDLTNGQLTPNNNTPLLAQALSVMNNYQAAAAGFGP
PASPHAAGGRAGFPAASSPGPALDVYGSAADSVGYVQAASPQPSGFPAIAVSRAPLNYTPGPLIPAAFTN
GYH


>gnl|WGS|Hap1_Pmyltmp_010581-P1 adult-specific cuticular protein ACP-20-like
MTLMIVLCFVLMMVAHLQAHEHAFSSNHISRHDGHHKEVHHEHDHHHDYYTHPKYEFEYKVEDHHTGDKK
SQHEHRDGDVVKGGYSFHQPDGSERHVEYHSDDHSGFHADVKYSTHHIVPHHHHY


>gnl|WGS|Hap2_Pmyltmp_010651-P1 uncharacterized protein Pmyltmp_010651
MAYTTIDTPVNSLSTMPFKPEHSNIRLVTVIRAATLKNIEDPRQKRPTTSNMIVLCFALMMVAHLQAHEH
AFSSNHISRHDGHHKEVHHEHDHHHDYYTHPKYEFEYKVEDHHTGDKKSQHEHRDGDVVKGGYSFHQPDG
SERHVEYHSDDHSGFHADVKYSTHHIVPKHHH


>gnl|WGS|Hap2_Pmyltmp_000911-P1 prostaglandin reductase 1-like isoform 1-T1
MVRARKYVVKNHFKGLPKPEDYEIVEYELPPLQHEEILIKTEWISVDPYHRVYNVYYQVPYDQFGFQVGT
VLESKNPKFPVGTKVVTHKGWCDYAIAKEGFAYDSGPIRHIYKLPNLEGLSQSLGVGAVGMPGVTAYFGF
LDICQPKPGETVVVTGAAGAVGSLVGQIAKIKGCRVIGFAGSDDKCQWLEKELGFDKAINYKTTEISKAL
KEAAPNGVDCYFDNVGGEVSSTIIYQMNKFGRVSVCGSISAYNEDVSKRPKASILQPPIVFNQLKIQGFV
ILEWVDRWQEAFDQLTKWIKSGQLIAREHVTDGFENVFDAFVGMLAGENTGKAVVKV
```
